# An Integrated Framework for Transcriptomic Characterization and Lorentzian Hyperbolic Visualization of a High-Risk Topological Branch in Alzheimer’s Disease

**DOI:** 10.64898/2026.06.13.732037

**Authors:** Chenhao Zeng, Zhibin Pu, Yuyang Tao, Wenshi Wei, Jian Zhao, Mingliang Cai, Shufei Ge

## Abstract

Alzheimer’s disease (AD) is a highly heterogeneous brain disorder in which molecular alterations vary across brain regions, disease stages, and patient subgroups. This study introduces an integrated analytical framework for characterizing transcriptomic variation associated with a high-risk topological branch, which was identified based on Lorentz distance in postmortem Brodmann area 36 samples from the Mount Sinai Brain Bank cohort, where over 70% of samples were in Braak stages V––VI. The framework integrates weighted gene co-expression network analysis, repeated stability-based differential expression analysis, network-level gene filtering, Gene Ontology enrichment, and nested stratified cross-validation to evaluate whether topological branch-associated genes capture biologically meaningful signals and carry predictive information for high-Braak group status. The identified gene sets were functionally enriched for neuronal development, neuron projection organization, synaptic signaling, vesicle fusion, and regulated synaptic release, suggesting that the high-risk topological branch reflects biologically relevant transcriptomic programs linked to neurodegenerative progression. Nested cross-validation further showed that the selected genes achieved measurable internal predictive performance for distinguishing high-Braak samples. As a second methodological contribution, we introduced a Lorentzian hyperbolic variant of t-distributed stochastic neighbor embedding (Lorentz t-SNE) to explore latent non-Euclidean structure in transcriptomic data. This method embeds samples in hyperbolic space, providing an alternative to Euclidean embeddings for representing hierarchical or nonlinear structures. Compared with conventional Euclidean embeddings, the proposed Lorentz t-SNE revealed a more localized organization of high-Braak samples. Together, these results demonstrate the utility of the proposed analytical framework and Lorentz t-SNE for investigating heterogeneous, potentially non-Euclidean organization in AD transcriptomes.

## 1 Introduction

Alzheimer’s disease (AD) is the most common cause of dementia and is pathologically characterized by extracellular amyloid-*β* deposition and intracellular tau neurofibrillary tangles [Scheltens et al., 2016, 2021]. The Braak staging framework characterizes a spatial propagation pattern of tau pathology in AD: tau-related pathology typically begins in the entorhinal cortex, subsequently involves limbic regions, and eventually extends to widespread neocortical areas in a relatively orderly manner [Braak and Braak, 1991, 1995, Braak et al., 2006]. This non-uniform distribution of pathology has motivated the concept of selective regional vulnerability, whereby distinct brain regions and cellular populations differ in their susceptibility to AD-related pathological processes [Roussarie et al., 2020]. Large-scale postmortem brain transcriptomic studies provide direct evidence for this theory: AD-associated gene expression changes are not uniformly distributed across the brain, but vary significantly with anatomical region, the burden of amyloid plaques and neurofibrillary tangles, and disease stage [Zhang et al., 2013, Wang et al., 2016, Wan et al., 2020]. Single-cell and single-nucleus transcriptomic studies have further demonstrated that AD-related molecular alterations are cell-type-specific and region-dependent, suggesting that disease-associated expression changes are shaped jointly by regional microenvironment, cellular composition, and cell-state variation [Mathys et al., 2019, 2024]. In addition, molecular subtyping and transcriptomic stratification analyses based on bulk RNA-seq have identified reproducible AD molecular subtypes and transcriptomic endophenotypes, supporting substantial molecular heterogeneity at the sample level [Milind et al., 2020, Neff et al., 2021]. These findings suggest that, beyond individual differentially expressed genes, pathways, or co-expression modules, AD-related transcriptomic data may contain higher-order sample-level organization that remains incompletely characterized.

Most computational analyses of AD transcriptomic data have been conducted within conventional vector-space or network-based frameworks, in which each sample is typically represented as a high-dimensional vector defined by its gene-expression profile, and downstream analyses primarily rely on gene-wise statistics, linear projection, correlation structure, or co-expression relationships. Differential expression analysis has been widely used to identify genes and pathways associated with disease status or pathological severity, including early microarray studies of AD brain tissue as well as subsequent RNA-seq and integrative transcriptomic analyses [Blalock et al., 2004, Ritchie et al., 2015, Zhang et al., 2013]. In parallel, weighted gene co-expression network analysis (WGCNA) and related network-based methods have provided an important framework for identifying co-regulated gene modules, hub genes, and disease-associated molecular programs in AD [Langfelder and Horvath, 2008, Zhang et al., 2013, Wang et al., 2016, Liang et al., 2018, Hu et al., 2020]. These approaches have substantially advanced the molecular characterization of AD and remain critical tools for identifying dysregulated genes, pathways, and co-expression modules. However, these methods may fail to fully preserve more complex latent structures within the data—an important consideration given that high-dimensional biological data can exhibit nonlinear transitions, branching architectures, and other manifold-like patterns, as revealed by manifold learning approaches in biological data visualization [Moon et al., 2019]. For a progressive and highly heterogeneous disease such as AD, the inherent organization of transcriptomic samples may therefore extend beyond simple linear separation or uniformly distributed variation. This motivates further investigation into the latent organizational structure of AD-related transcriptomic samples and its appropriate geometric representation.

Hyperbolic geometry provides a non-Euclidean framework for modeling latent structures that are poorly captured by Euclidean space. Owing to its constant negative curvature and exponential volume growth, it is well suited for representing hierarchical or tree-like relationships with low distortion [Nickel and Kiela, 2017, 2018]. These properties have motivated the application of hyperbolic representation learning to complex biological data. In transcriptomics, low-dimensional hyperbolic geometry has been shown to characterize multiple gene-expression datasets and to reveal latent expression structures that may be obscured in Euclidean visualizations [Zhou and Sharpee, 2021]. In single-cell genomics, Poincaré maps embed scRNA-seq data in the Poincaré disk to visualize cellular hierarchies, lineage relationships, and pseudotemporal organization [Klimovskaia et al., 2020]. Subsequent methods have extended non-Euclidean modeling through probabilistic and deep-learning frameworks, including scPhere for hyperspherical or hyperbolic latent representations of scRNA-seq data [Ding and Regev, 2021], scDHMap for hyperbolic manifold learning of sparse single-cell count data [Tian et al., 2023], and PoincaréDMT for modeling branching trajectories, batch effects, and marker-gene structure [Xu et al., 2025]. Recent extensions to spatial transcriptomics include MManiST, which integrates Euclidean and hyperbolic manifold encoders for spatial domain identification [Li et al., 2025], and HyperDiffuseNet, which combines hyperbolic latent modeling with graph diffusion for dimensionality reduction and representation learning [Qi et al., 2025]. Hyperbolic neural architectures have also been applied to genome sequence representation learning [Khan et al., 2025]. Collectively, these studies suggest that hyperbolic representations provide a useful framework for biological data whose latent organization is poorly captured by Euclidean geometry.

In the topological structure analysis of the Mount Sinai Brain Bank (MSBB) cohort, the Gaussian mixture model (GMM) soft Mapper algorithm generated the topological structure of transcriptomic samples from Brodmann area 36 (corresponding to the parahippocampal gyrus), and one branch was identified as a high-risk topological branch and exhibited a clear association with AD severity patterns [Tao and Ge, 2025]. However, the biological and geometric interpretation of this branch remains incomplete. In particular, it remains unclear how this topology-derived branch relates to Braak-defined pathological grouping, which co-expression modules and hub genes are associated with branch membership, and whether a stable set of branch-associated differentially expressed genes can be identified. In addition, because this branch was derived from a topology-based representation of sample organization involving a Lorentz distance structure, it is necessary to examine whether its organization can be further explored using a geometrically consistent visualization framework rather than only conventional Euclidean embeddings. These questions motivate an integrated analysis that links topology-derived branch membership with transcriptomic network structure, robust gene-level signatures, predictive evaluation, and hyperbolic visualization.

In this study, inspired by the high-risk topological branch reported in the literature [Tao and Ge, 2025], we conducted an in-depth analysis of the cerebral cortical Brodmann area 36, aiming to investigate the characteristics of the brain transcriptomic architecture associated with AD. We focused on two complementary sample annotations: membership in the Mapper-derived high-risk branch and Braak-defined high-pathology status. We first applied weighted gene co-expression network analysis to identify modules and hub genes associated with these annotations, and then used repeated stability-based differential expression analysis to select genes consistently associated with the high-risk branch. Functional enrichment analysis was performed to characterize the biological processes represented at the module, hub-gene, and stable-differential-expression levels. To assess whether the resulting branch-associated gene panel carries information relevant to AD pathology, we evaluated its ability to predict high-Braak group status using nested cross-validation. Finally, we applied the proposed Lorentz t-SNE to visualize sample organization under the Lorentz distance structure and compared it with the standard Euclidean t-SNE as a qualitative reference. Through this workflow, summarized in Figure 1, we aimed to characterize the transcriptomic features of the identified high-risk topological branch, evaluate its association with high-Braak group status through predictive modeling, and explore whether the Lorentz-based hyperbolic visualization can reveal aspects of sample organization that are less apparent under conventional Euclidean visualization.

**Figure 1:**
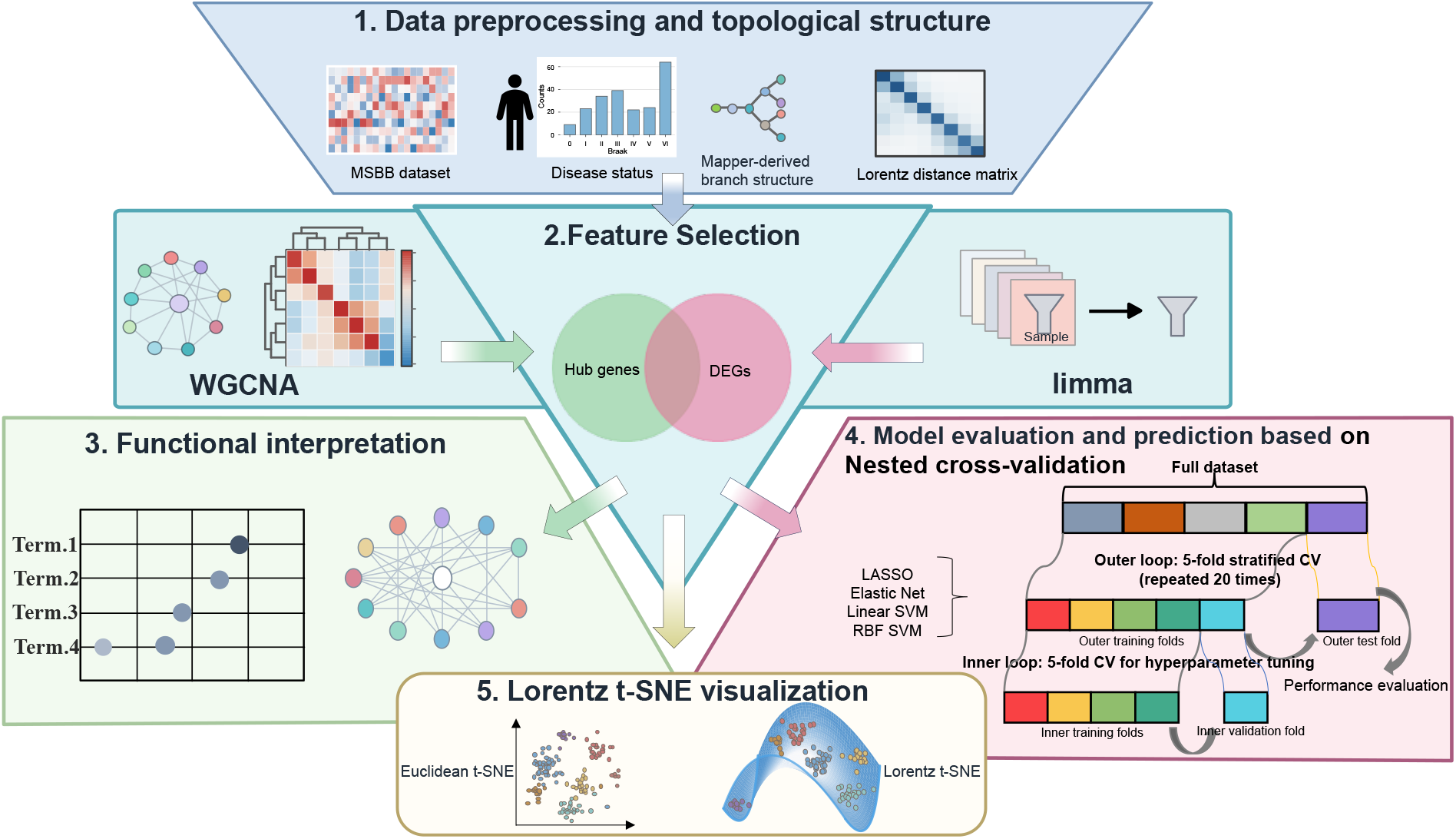
Overview of the study design and analysis workflow. The study began with MSBB transcriptomic data, and the GMM soft Mapper-derived topology-based branch membership. Feature selection integrated WGCNA-based hub genes with stable differentially expressed genes identified through repeated limma analyses. Branch-associated gene sets were then examined by functional enrichment analysis. A candidate gene panel was evaluated for prediction of high-Braak group status using nested cross-validation. Finally, the Lorentz t-SNE was used to visualize sample organization under the Lorentz distance structure and was compared with the standard Euclidean t-SNE to assess differences in the resulting sample layouts.

## 2 Materials and methods

### 2.1 Data collection and preprocessing

The data used in this study were obtained from the MSBB cohort through the AD Knowledge Portal [Wang et al., 2018]. We used the processed gene-level expression matrix provided by the AD Knowledge Portal. According to the MSBB RNA-seq processing pipeline, gene-level raw counts were converted to counts per million (CPM), normalized using the trimmed mean of M-values (TMM) method to account for differences in library size, and adjusted for known technical and biological covariates. These covariates included sequencing batch, sex, race, age, RNA integrity number (RIN), post-mortem interval (PMI), exonic mapping rate, and rRNA rate. The resulting covariate-adjusted log-scale expression matrix was used for all downstream analyses.

We applied the GMM soft Mapper algorithm to four MSBB brain regions, including the frontal pole in area 10 (FP; BM10), the superior temporal gyrus in area 22 (STG; BM22), the parahippocampal gyrus in area 36 (PHG; BM36), and the inferior frontal gyrus in area 44 (IFG; BM44) [Wang et al., 2018], and adopted the same data preprocessing strategy as in [Tao and Ge, 2025]. The complete output results of the GMM soft Mapper algorithm across these four regions are provided in Appendix Figure 9. The results revealed a distinct branched topological structure in the BM36 region’s GMM soft Mapper output. Over 70% of individuals located on this branch had high Braak scores (e.g., Braak stages V–VI), and it was referred to as the high-risk topological branch (see red dashed box in Appendix Figure 9a).

Such a pattern was not observed in the other three brain regions. Therefore, in this study, we focused on BM36 to evaluate the transcriptomic, functional, predictive, and geometric relevance of this topology-derived branch. In addition, the BM36 is located within medial temporal and parahippocampal regions that are relevant to AD-related pathology [Braak and Braak, 1991, Braak et al., 2006, Berron et al., 2021]. Using brain region BM36 allowed us to evaluate the transcriptomic and predictive relevance of this previously defined topological branch.

For the BM36 dataset, a total of 215 postmortem human brain tissue samples with annotations were included, each with over 20,000 gene expression measurements and an assigned Braak staging score ranging from 0 to VI, with higher scores indicating more advanced Alzheimer’s disease-related neuropathological stage [Braak and Braak, 1991, Hyman, 1998]. Samples were characterized using two complementary classification schemes. First, according to Braak staging, samples were classified into the high-Braak group (Braak stages V–VI; 88 samples) and the non-high-Braak group (Braak 0–IV; 127 samples). Second, based on branch assignment defined by the GMM soft Mapper analysis, we divided the samples into high-risk topological branch samples (22 cases) and samples from other branches (193 cases). The result aligns with prior findings [Tao and Ge, 2025]. Notably, among the 22 samples in the high-risk topological branch, 16 (73%) were concurrently classified into the high-Braak group, whereas the proportion of high-Braak group in the rest branches was only 37%. This result suggests a correspondence between molecular subgroups defined by topological structure and pathological groups defined by Braak staging. The overall distribution of Braak staging scores in the entire MSBB BM36 cohort, along with the distribution stratified by the high-risk topological branch versus all other branches, are shown in Appendix Figure 7a, b.

### 2.2 Weighted gene co-expression network analysis

Weighted Gene Co-expression Network Analysis (WGCNA) is a classic method for identifying phenotype-associated gene modules: it constructs a weighted correlation network between genes, clusters genes with similar expression patterns into functional modules, and analyzes gene synergy [Zhang et al., 2005, Langfelder and Horvath, 2008]. This study used WGCNA to identify co-expression modules associated with the high-risk topological branch. This strategy not only provides network-level evidence for screening branch-specific genes but also verifies the association between modules and high Braak stages (representing severe tau pathological burden), thereby providing network-level functional context for branch-associated transcriptional variation. To construct the co-expression network, genes were ranked according to their median absolute deviation (MAD), and the top 25% most variable genes were retained to reduce noise and computational complexity.

A signed co-expression network was constructed using biweight mid-correlation (bicor) to improve robustness to outlying expression values. The soft-thresholding power (*β*) was selected according to the scale-free topology criterion, and the adjacency matrix was transformed into a signed topological overlap matrix (TOM). Modules were identified using a signed network configuration, and highly correlated modules were subsequently merged based on eigengene distance.

Module eigengenes (MEs) were calculated to summarize the expression profile of each module. Each ME was then correlated with branch membership and Braak-defined group status to assess module-level associations with the topology-derived branch and AD-related pathological grouping. Statistical significance for module–trait associations was adjusted using the Benjamini–Hochberg procedure to control the false discovery rate (FDR) [Benjamini and Hochberg, 1995]. For downstream hub-gene prioritization, we focused on the modules showing the strongest absolute correlations with branch membership. Within these branch-associated modules, hub genes were defined using module membership (*MM*, |*MM* | ≥ 0.60) and branch-based gene significance (*GS*_branch_, |*GS*_branch_| ≥ 0.25).

### 2.3 Differential gene expression analysis and feature selection

Given the modest cohort size and the imbalance in branch membership, differential expression results based on a single data split may be sensitive to the allocation of samples across folds, particularly for the high-risk topological branch. We therefore used a repeated resampling-based differential expression strategy to select genes showing consistent branch-associated expression differences rather than relying on a single full-cohort comparison. Specifically, 100 resampled training subsets were generated using 20 repetitions of stratified 5-fold partitioning, following a repeated partitioning scheme consistent with the downstream cross-validation framework. Within each subset, differential expression analysis was performed using `limma` by comparing samples in the high-risk topological branch with samples from the other branches [Smyth, 2004, Ritchie et al., 2015]. Differentially expressed genes were identified using predefined criteria of FDR < 0.05 and | log_2_ FC| > 1.

For each gene, the selection frequency was defined as the number of resampled differential expression analyses in which the gene satisfied the predefined significance and effect-size criteria. Genes selected in at least 80 of the 100 iterations were defined as stable differentially expressed genes (stable DEGs) associated with the high-risk topological branch. This frequency-based criterion was used to retain genes with reproducible branch-associated expression differences across repeated resampling while reducing dependence on any single data partition, following the general rationale of stability selection [Meinshausen and Bühlmann, 2010].

To incorporate network-level information, these stable DEGs were intersected with WGCNA-derived hub genes from the branch-associated modules. Because both the differential expression comparison and the hub-gene selection were defined with respect to branch membership rather than high-Braak group status, the resulting candidate panel was treated as a branch-associated gene set. This fixed candidate panel was subsequently used for functional enrichment analysis and for internal predictive evaluation of high-Braak group status.

### 2.4 Functional enrichment analysis

To provide functional context for the branch-associated gene sets identified at different analytical layers, Gene Ontology (GO) Biological Process (BP) enrichment analysis [Ashburner et al., 2000] was performed for branch-associated module genes, WGCNA-derived hub genes, and stable differentially expressed genes (DEGs). Enrichment analysis was conducted in R using the `clusterProfiler` package [Yu et al., 2012, Wu et al., 2021]. Gene identifiers were supplied to `clusterProfiler` as Ensembl gene IDs, corresponding to the ENSG-formatted identifiers in the processed expression matrix.

The background gene set for each enrichment analysis was defined according to the gene space from which the corresponding target gene set was selected. In this study, the WGCNA input genes were used as the background gene set for the module-level and hub-gene analyses, whereas the genes included in the repeated differential expression analysis were used as the background gene set for stable DEGs. This analysis-specific background definition was used to reduce potential enrichment bias caused by comparing target gene sets against an inappropriate or overly broad background gene set [Timmons et al., 2015]. GO-BP enrichment was used as the primary functional annotation framework because it provides interpretable biological process-level summaries of branch-associated transcriptional signals.

Redundant GO terms were reduced, where appropriate, based on semantic similarity using semantic-similarity-based filtering supported by GOSemSim [Yu et al., 2010]. Statistical significance was assessed using Benjamini–Hochberg false discovery rate (FDR) correction [Benjamini and Hochberg, 1995], and terms with adjusted *P* values < 0.05 were considered significant. Enrichment results were visualized according to the corresponding analytical layer, including a dotplot for module-level enrichment, an enrichment map for hub-gene enrichment, and a horizontal barplot for stable-DEG enrichment. These visualizations were used to summarize representative enriched GO-BP terms, whereas the full enrichment results were retained for supplementary reporting.

### 2.5 Nested cross-validation-based prediction modeling and evaluation of high-Braak group status

To evaluate whether the branch-associated candidate genes contained information relevant to the prediction of high-Braak group status, we used a nested repeated stratified 5-fold cross-validation framework for model training, hyperparameter tuning, and performance evaluation [Varma and Simon, 2006]. Because of the modest cohort size and the imbalanced branch membership, feature discovery from a single data partition may depend on how samples are allocated, especially for the small high-risk topological branch. To reduce this dependence and to avoid re-selecting features using information from held-out test folds during model evaluation, the branch-associated candidate gene panel was defined before cross-validation modeling based on the stability-based differential expression analysis and WGCNA filtering strategy described above. This predefined candidate gene panel was then used as the input feature set throughout the nested cross-validation procedure, providing internal performance estimates for this fixed feature set while limiting selection bias and overly optimistic performance estimates in high-dimensional predictive modeling [Ambroise and McLachlan, 2002, Cawley and Talbot, 2010, Varma and Simon, 2006].

To ensure that each cross-validation fold contained a reasonable proportion of samples from both the high-Braak group and the high-risk topological branch, we constructed a joint stratification variable based on the combination of these two labels and used it to generate the cross-validation splits. The outer 5-fold cross-validation procedure was repeated 20 times with different random seeds to obtain more stable estimates of model performance. In each outer iteration, one fold was held out exclusively for testing, whereas the remaining folds were used for model fitting and inner-loop hyperparameter tuning. The held-out test fold was not used during model selection. Within each outer-loop training fold, class weights were calculated using only the samples in that training fold to account for class imbalance. Feature scaling, including mean centering and variance scaling, was performed only on the outer-loop training data, and the same transformation was then applied to the corresponding held-out test fold to avoid information leakage.

Within each outer-loop training fold, an independent inner 5-fold cross-validation procedure was used for hyperparameter tuning. Four classifiers were evaluated: logistic regression with Lasso regularization, elastic net logistic regression, linear support vector machine (SVM), and radial basis function (RBF) SVM. For the SVM models, grid search was used to select the cost parameter *C* and, for the RBF kernel, the kernel parameter *γ*, with model selection based on the mean PR-AUC across inner validation folds. For the penalized logistic regression models, the regularization parameter *λ* was selected using the 1-standard-error rule within the inner cross-validation procedure [Friedman et al., 2010].

For each outer fold, the model configuration selected in the inner loop was retrained on the full outer-loop training fold and evaluated on the held-out test fold. Predictive performance was assessed using ROC-AUC and PR-AUC, together with sensitivity, specificity, and Matthews correlation coefficient (MCC) [Matthews, 1975]. Because the high-Braak classification task involved class imbalance, PR-AUC was used as a complementary threshold-independent metric alongside ROC-AUC [Davis and Goadrich, 2006, Saito and Rehmsmeier, 2015]. A probability threshold of 0.5 was used to derive threshold-dependent metrics, including sensitivity, specificity, and MCC.

After nested cross-validation, the classifier with the most favorable internal performance profile was used as the primary model for interpreting gene contributions within the candidate panel. This primary model was refit on the full cohort to estimate final model coefficients; for regularized logistic regression, the regularization parameter was selected by internal 5-fold cross-validation using the 1-standard-error (1-SE) rule. Genes assigned non-zero coefficients in the refitted model were retained for subsequent model interpretation. The refitted model can be expressed as a linear predictor formed by the weighted standardized expression values of these genes. Model performance was evaluated using predicted probabilities from held-out outer folds generated during nested cross-validation; the full-cohort refitted model was used only to estimate coefficients, display non-zero coefficient genes, and interpret the model structure.

### 2.6 Lorentz t-SNE

To further examine the sample organization associated with the Mapper-derived high-risk topological branch, we visualized the MSBB BM36 transcriptomic samples using both the standard Euclidean t-SNE and our proposed Lorentz-hyperbolic variant of t-SNE. The Mapper-based analysis of the MSBB dataset used implicit intervals to identify a branching sample organization and derived a sample-wise Lorentz distance matrix to characterize inter-sample dissimilarities [Tao and Ge, 2025]. This motivated us to examine whether a low-dimensional hyperbolic embedding could provide an additional visualization of the same distance structure.

For the Euclidean reference visualization, the standard t-SNE was applied using the sample-wise Lorentz distance matrix as a precomputed dissimilarity measure [Maaten and Hinton, 2008]. As an additional reference analysis, we also applied standard Euclidean t-SNE using the Euclidean distance matrix computed from the processed BM36 expression profiles. For the Lorentz t-SNE, the low-dimensional embedding was defined on the two-dimensional Lorentz hyperboloid ℋ^2^, which is embedded in ℝ^3^. The choice of *n* = 2 was made to obtain a two-dimensional hyperbolic representation suitable for visualization. Low-dimensional similarities were computed from Lorentz geodesic distances on ℋ^2^, and the embedding was optimized by minimizing the Kullback–Leibler divergence using Riemannian gradient descent [Absil et al., 2008, Bonnabel, 2013]. For visualization, the optimized Lorentz-hyperboloid coordinates were mapped to their corresponding two-dimensional Poincaré disk coordinates.

The Lorentz model was used rather than optimizing directly in the Poincaré disk for two reasons. First, the branch structure examined in this study was originally derived from a Lorentz distance-based analysis, and using the Lorentz model allowed the visualization procedure to remain consistent with the corresponding dissimilarity structure. Second, the Lorentz model has been widely used for hyperbolic representation learning and has favorable optimization properties relative to the Poincaré ball in several hyperbolic embedding settings [Nickel and Kiela, 2018, Chami et al., 2019, Mishne et al., 2023]. The algorithmic procedure used in this study is outlined in Algorithm 1, and the mathematical details of the Lorentz hyperboloid model, Riemannian gradient, and exponential-map update are provided in Appendix 1.

## 3 Results

### 3.1 WGCNA

We performed WGCNA to identify co-expression modules associated with the high-risk topological branch and to examine their correspondence with high-Braak group status. Based on the scale-free topology criterion, the soft-thresholding power was set to *β* = 10, at which the scale-free topology fit index exceeded 0.85 while maintaining adequate mean connectivity (Figure 2a, b).

**Figure 2:**
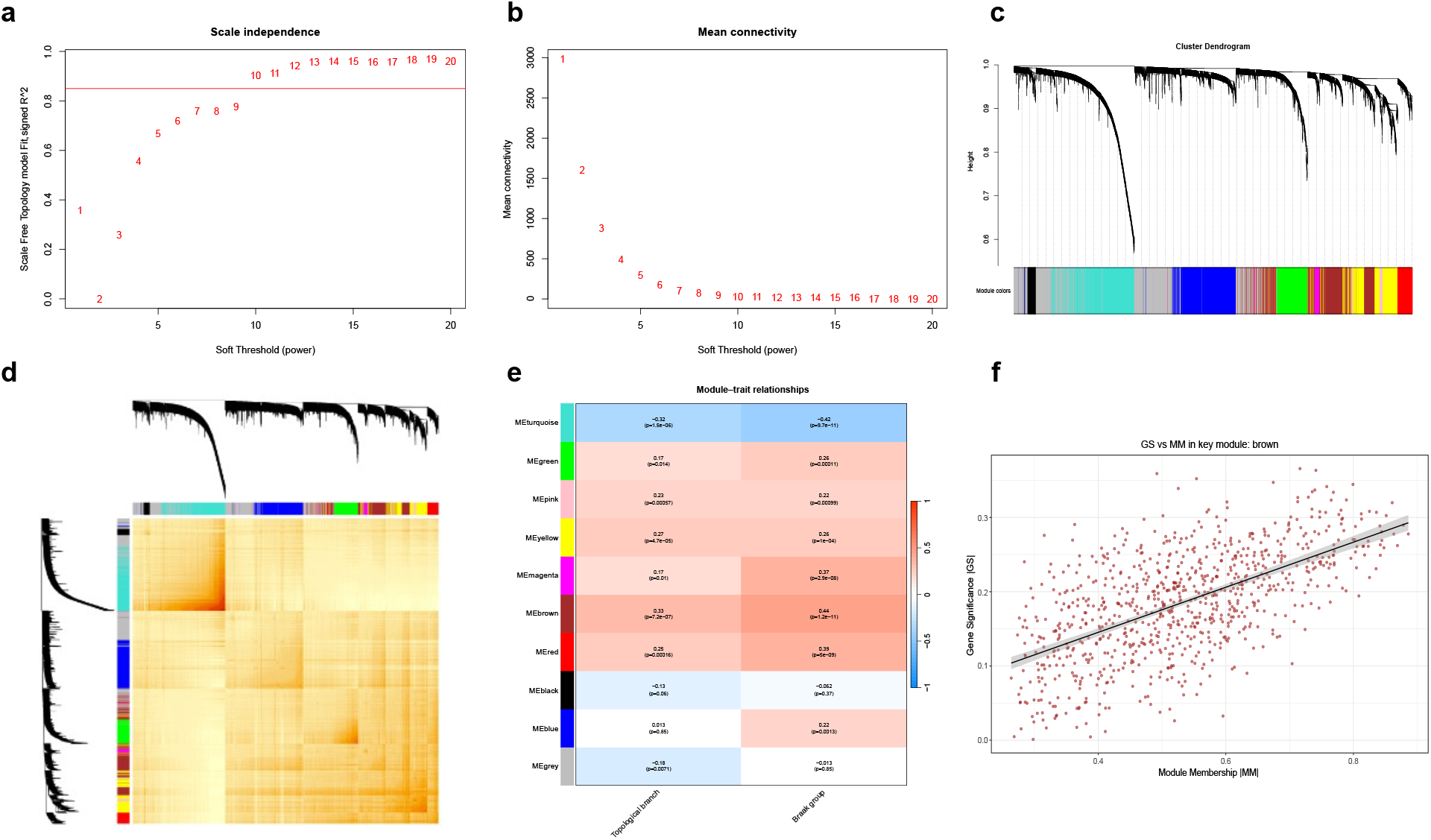
Weighted gene co-expression network analysis (WGCNA) identifying modules associated with the high-risk topological branch and the high-Braak group (Braak V–VI). **(a)** Scale-free topology fit index for different soft-thresholding powers. **(b)** Mean connectivity for different soft-thresholding powers. **(c)** Hierarchical clustering dendrogram of genes based on topological overlap dissimilarity. The color bar below the dendrogram indicates the merged WGCNA module assignment of each gene, with each color representing one co-expression module. **(d)** Heatmap of the topological overlap matrix (TOM). Rows and columns represent genes ordered according to the clustering dendrogram, and the module color bars along the top and left margins indicate the same merged WGCNA module assignments as in panel c. Color intensity within the heatmap represents topological overlap between gene pairs, and block-like regions reflect modular structure in the co-expression network. **(e)** Heatmap of correlations between module eigengenes and the study traits, including high-risk topological branch membership and high-Braak group status (Braak V–VI). Rows correspond to module eigengenes, and the module labels/colors correspond to the same WGCNA modules shown in panels c and d. Heatmap colors represent correlation coefficients, with red indicating positive correlations and blue indicating negative correlations. Numbers outside parentheses indicate correlation coefficients, and the corresponding *P* -values are shown in parentheses. **(f)** Relationship between module membership and gene significance in the brown module.

Genes were subsequently grouped into co-expression modules based on topological overlap, and modules with similar eigengene profiles were merged (Figure 2c). The TOM heatmap showed a block-like pattern, consistent with the presence of modular structure in the constructed co-expression network (Figure 2d).

Correlation analysis between module eigengenes and the study traits showed that the **brown** and **turquoise** modules had the strongest absolute correlations with high-risk topological branch membership (Figure 2e). The brown module was positively correlated with the high-risk topological branch (*r* = 0.33, *P* = 7.2 × 10^−7^), whereas the turquoise module was negatively correlated with branch membership (*r* = −0.32, *P* = 1.5 × 10^−6^). Notably, these two branch-associated modules also showed strong associations with high-Braak group status, with the brown module positively correlated with the high-Braak group (*r* = 0.44, *P* = 1.2 × 10^−11^) and the turquoise module negatively correlated with the high-Braak group (*r* = −0.42, *P* = 9.7 × 10^−11^).

We next examined the relationship between module membership and branch-based gene significance within the branch-associated modules. In the brown module, genes with higher module membership tended to show stronger associations with branch membership, indicating concordance between intramodular connectivity and branch-based gene significance (Figure 2f). Based on the predefined thresholds of |*MM*| ≥ 0.60 and | *GS*_branch_ | ≥ 0.25, hub genes from the brown and turquoise modules were retained for subsequent analyses, resulting in a total of 305 branch-associated hub genes.

(The WGCNA-related supplementary results are provided as Supplementary Data: Supplementary Data 1 contains the MAD-filtered WGCNA input genes, and Supplementary Data 2 contains the module–trait association results together with the WGCNA-derived branch-associated hub genes.)

### 3.2 Identification of stable differentially expressed genes associated with the high-risk topological branch

We first performed a conventional differential expression analysis comparing samples in the high-risk topological branch with samples from the other branches. The resulting volcano plot showed both upregulated and downregulated genes, indicating detectable branch-associated transcriptional differences under the predefined significance and effect-size criteria (Figure 3a). Given the high dimensionality of the expression matrix, the modest number of samples in the high-risk branch, and the potential sensitivity of single-run differential expression analysis to sample partitioning, we next applied a repeated resampling-based strategy to identify genes that were consistently selected across iterations.

**Figure 3:**
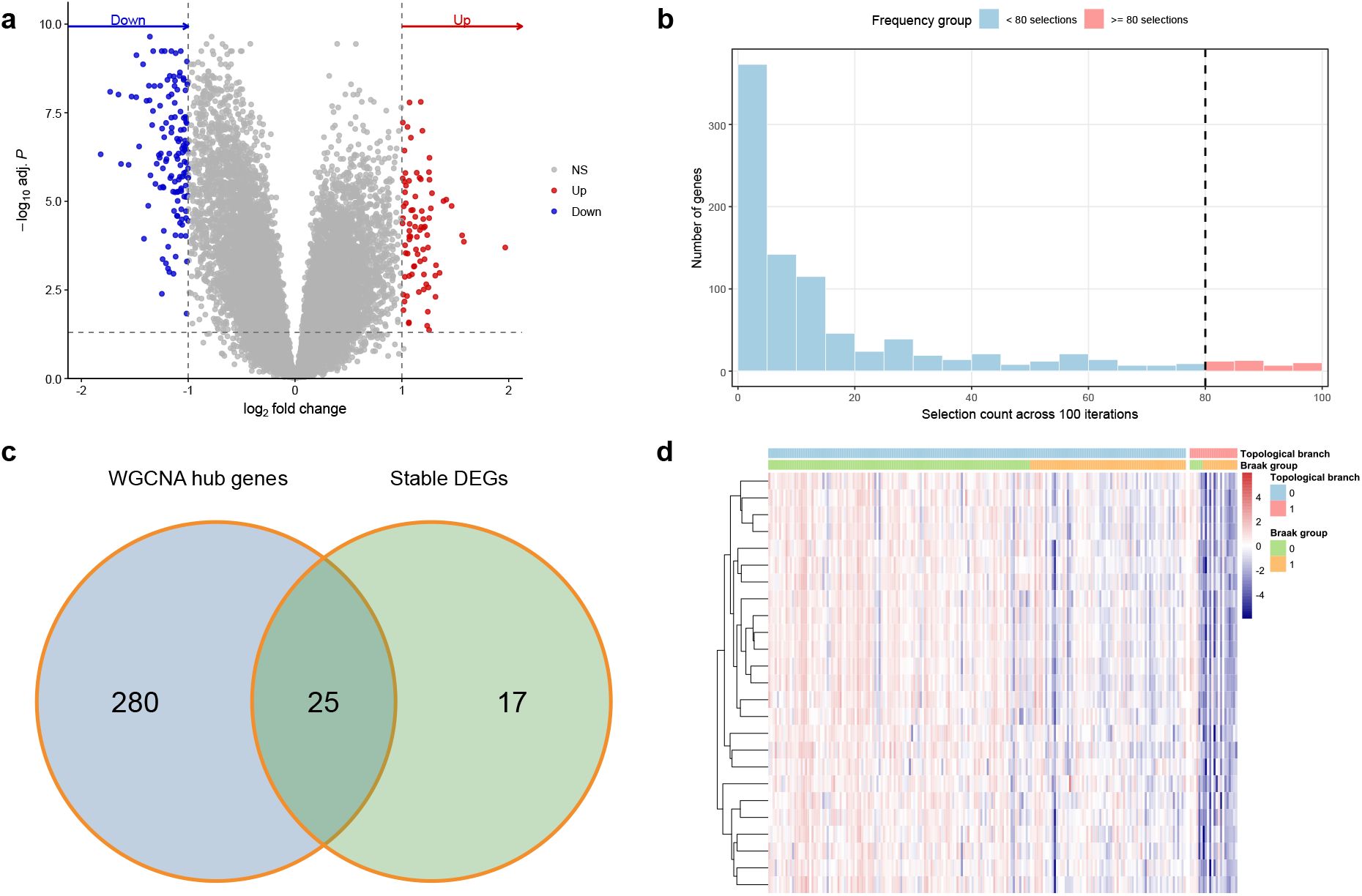
Identification of stable differentially expressed genes associated with the high-risk topological branch. **(a)** Volcano plot from the conventional differential expression analysis comparing the high-risk topological branch with samples from the other branches. Red and blue dots indicate significantly upregulated and downregulated genes, respectively. **(b)** Distribution of gene selection frequencies derived from repeated limma-based differential expression analyses across 100 resampled training subsets generated from a 20-repeat 5-fold partitioning scheme. The black dashed line indicates the predefined stability threshold, and the red bars highlight genes retained in at least 80 of the 100 iterations. **(c)** Intersection of WGCNA hub genes and stable DEGs, resulting in a 25-gene branch-associated candidate panel for downstream analyses. **(d)** Heatmap of the 25-gene branch-associated candidate panel across BM36 samples. Color indicates gene-wise standardized expression, with red indicating relatively higher expression and blue indicating relatively lower expression. The upper annotation bars indicate topological branch membership and Braak group status; topological branch 1 denotes the high-risk topological branch and topological branch 0 denotes samples from the other branches, whereas Braak group 1 denotes high-Braak samples and Braak group 0 denotes non-high-Braak samples.

Specifically, `limma` analysis was repeated across 100 resampled training subsets, and the selection frequency of each gene was defined as the number of iterations in which it satisfied the predefined differential expression criteria. The resulting frequency distribution showed that many genes were selected only occasionally, whereas a smaller subset was repeatedly retained across resampling iterations (Figure 3b). This pattern supported the use of a frequency-based threshold to distinguish consistently selected branch-associated genes from genes whose selection was more dependent on individual data splits.

Using the predefined stability threshold, genes selected in at least 80 of the 100 iterations were retained as stable differentially expressed genes associated with the high-risk topological branch. This procedure identified 42 stable DEGs. To incorporate network-level information, these stable DEGs were intersected with WGCNA-derived hub genes from the branch-associated brown and turquoise modules, resulting in a 25-gene candidate panel for downstream analyses (Figure 3c).

The heatmap of the 25-gene candidate panel showed a branch-related expression pattern. After samples were grouped by topological branch membership and ordered by Braak stage within each branch group, samples in the high-risk topological branch tended to show lower standardized expression across many candidate genes, whereas samples from the other branches displayed more heterogeneous expression patterns (Figure 3d). Taken together, these results indicate that repeated resampling-based differential expression analysis, combined with branch-associated network filtering, identified a set of candidate genes showing stable branch-associated expression differences, which was subsequently used for functional and predictive analyses. The gene selection-frequency results from the repeated resampling-based `limma` analyses, which followed a partitioning scheme consistent with the downstream nested cross-validation framework, together with the retained stable DEGs, are provided in Supplementary Data 3.

### 3.3 Functional enrichment analysis of branch-associated gene sets

To characterize the functional context of branch-associated transcriptional signals, GO-BP enrichment analyses were performed at the module, hub-gene, and stable-DEG levels. At the module level, branch-associated module genes were enriched for terms related to nervous system development, generation of neurons, synaptic signaling, chemical synaptic transmission, synaptic vesicle cycle, vesicle-mediated transport in synapse, and neurotransmitter secretion (Figure 4a). These results indicate that the broader branch-associated module landscape was linked to neuronal and synaptic biological processes.

**Figure 4:**
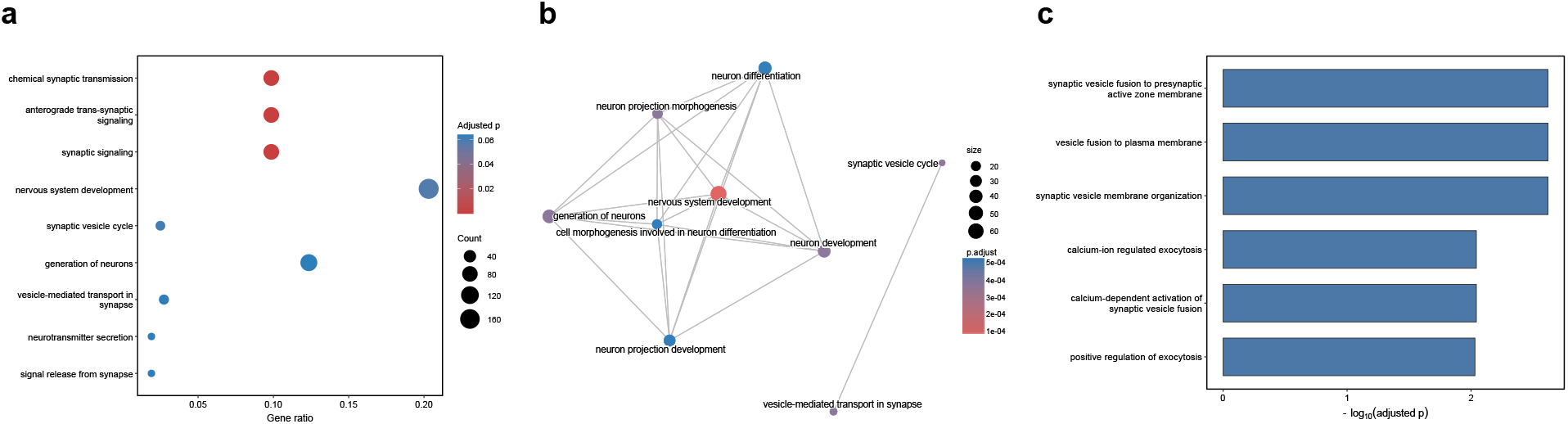
Gene Ontology Biological Process enrichment analysis of branch-associated gene sets. Adjusted *P* values shown in all panels were calculated using the Benjamini–Hochberg procedure. **(a)** Dotplot of representative enriched GO Biological Process terms for branch-associated module genes. The x-axis indicates the gene ratio, defined as the proportion of input genes annotated to each term. Dot size represents the number of input genes mapped to the corresponding term, and dot color represents the adjusted *P* value. **(b)** Enrichment map of representative enriched GO Biological Process terms for WGCNA-derived hub genes. Each node represents an enriched GO Biological Process term. Node size indicates the number of mapped genes, node color represents the adjusted *P* value, and edges indicate pairwise similarity between terms based on gene overlap or semantic relatedness. **(c)** Horizontal barplot of representative enriched GO Biological Process terms for stable differentially expressed genes. Bar length represents − log_10_-transformed adjusted *P* value.

When the analysis was restricted to WGCNA-derived hub genes, enriched terms were concentrated in processes related to nervous system development, neuron development, neuron differentiation, neuron projection development, and neuron projection morphogenesis. The enrichment map further showed that these terms formed an interconnected functional cluster rather than isolated signals (Figure 4b). This pattern suggests that the hub-gene layer captured a more focused subset of neuronal differentiation and projection-related processes within the broader branch-associated modules.

Stable DEGs identified through repeated `limma`-based selection showed enrichment for terms related to presynaptic and vesicle-associated processes, including synaptic vesicle fusion to the presynaptic active zone membrane, vesicle fusion to the plasma membrane, synaptic vesicle membrane organization, and calcium-regulated exocytosis (Figure 4c). Compared with the broader module- and hub-level enrichment patterns, the stable-DEG layer highlighted a smaller set of processes related to vesicle fusion and regulated synaptic release. These enrichment results should be interpreted as functional annotations of branch-associated gene sets rather than direct evidence of altered cellular mechanisms.

### 3.4 Model evaluation and prediction based on nested cross-validation

To evaluate whether the branch-associated candidate panel contained information relevant to the prediction of high-Braak group status, we compared four candidate classifiers within the nested cross-validation framework, including Logistic LASSO, Elastic Net Logistic, Linear SVM, and RBF SVM. Overall, all models showed moderate internal predictive performance, with the penalized logistic regression models showing slightly better overall performance and relatively greater stability across repeated outer-fold evaluations (Table 1). Among the evaluated classifiers, the Elastic Net Logistic model achieved the highest mean PR-AUC (0.759 ± 0.078) and a mean ROC-AUC of 0.789 ±0.065, together with the highest mean MCC (0.522 ± 0.124). The ROC and precision–recall curves based on aggregated out-of-fold (OOF) predicted probabilities were broadly consistent with this overall performance pattern (Figure 5a, b). The outer-fold performance distributions further suggested that model performance was relatively stable across repeated data splits, although the distributions of candidate models partially overlapped (Figure 5d).

**Table 1:**
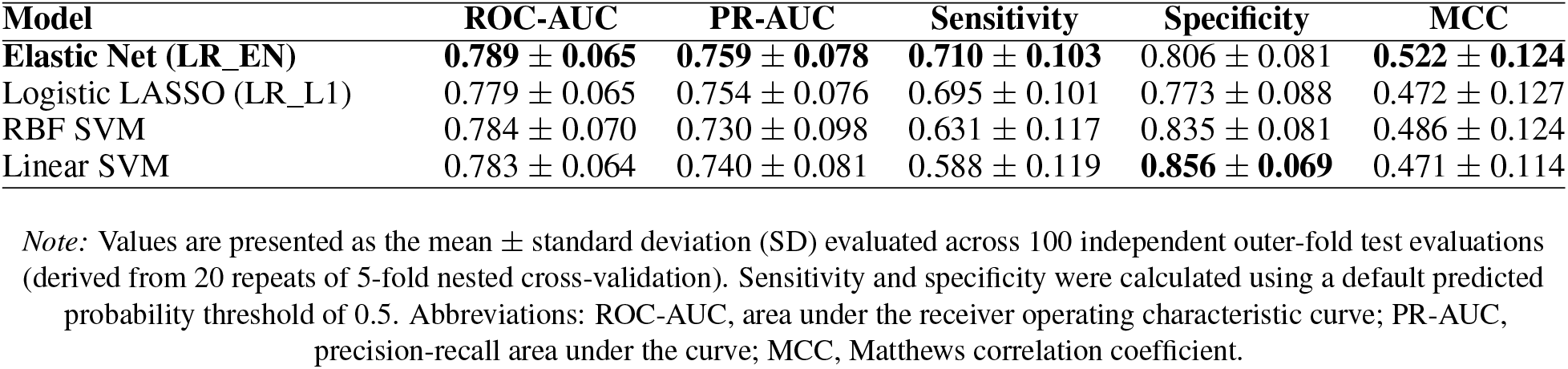
Performance summary of candidate models for prediction of high-Braak group status under nested cross-validation.

**Figure 5:**
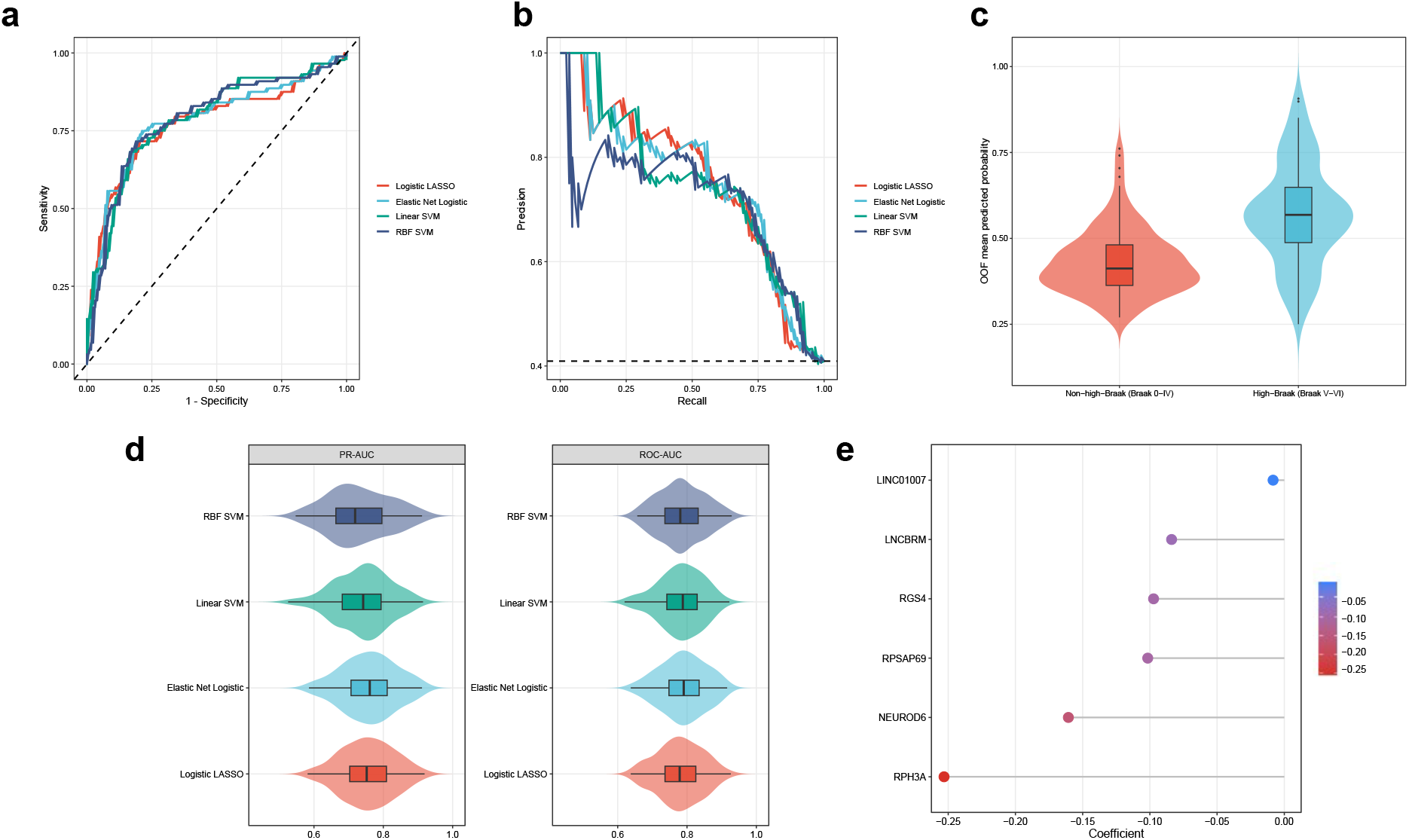
Model evaluation and prediction of high-Braak group status based on nested cross-validation. **(a)** ROC curves of the candidate models based on pooled OOF predictions. **(b)** Precision–recall curves of the candidate models based on pooled OOF predictions. **(c)** Distribution of OOF predicted probabilities in the non-high-Braak group and high-Braak group. **(d)** Distribution of PR-AUC and ROC-AUC across repeated outer-fold evaluations for each candidate model. **(e)** Coefficients of the six retained genes in the final Elastic Net model.

Given its overall internal performance, stability, and interpretability, the Elastic Net Logistic model was selected as the primary model for interpreting gene contributions within the candidate panel. The model was refit on the full estimate final coefficients. Under the combined *L*_1_*/L*_2_ penalization, only six genes from the 25-gene candidate panel retained non-zero coefficients in the refitted model (Figure 5e). All six retained genes showed negative coefficients, indicating that lower standardized expression of these genes contributed to a higher predicted probability of high-Braak group membership in the fitted model. This direction was broadly consistent with the branch-related expression pattern observed for the candidate panel.

For interpretability, the refitted Elastic Net logistic regression model was represented using the logit link function, Logit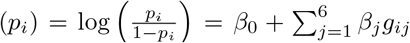, where *p*_*i*_ denotes the predicted probability that sample *i* belongs to the high-Braak group, *g*_*ij*_ denotes the standardized expression value of the *j*th retained gene in sample *i, β*_0_ denotes the intercept, and *β*_*j*_ denotes the corresponding non-zero Elastic Net coefficient. To facilitate reproducibility, the exact coefficients and the scaling parameters required for standardization are provided in Appendix Table 2. Model performance was evaluated using predicted probabilities generated from held-out outer folds during nested cross-validation, rather than predictions from the full-cohort refitted model.

**Table 2:** Coefficients and feature-scaling parameters of the genes retained in the full-cohort refitted Elastic Net logistic regression model.

| Index | Gene ID | Gene symbol | Feature type | Coefficient | Mean for scaling | SD for scaling |
| --- | --- | --- | --- | --- | --- | --- |
| – | – | Intercept | Intercept | -0.04344 | – | – |
| $\beta_1$ | ENSG00000089169 | RPH3A | Gene | -0.25311 | 6.642763 | 0.774701 |
| $\beta_2$ | ENSG00000164600 | NEUROD6 | Gene | -0.16062 | 4.256257 | 1.175136 |
| $\beta_3$ | ENSG00000233487 | RPSAP69 | Gene | -0.10160 | 1.563167 | 0.786669 |
| $\beta_4$ | ENSG00000117152 | RGS4 | Gene | -0.09726 | 8.262172 | 1.007956 |
| $\beta_5$ | ENSG00000249436 | LNCBRM | Gene | -0.08385 | 2.216202 | 1.000173 |
| $\beta_6$ | ENSG00000233123 | LINC01007 | Gene | -0.00836 | 1.983835 | 1.153468 |

Finally, we examined the distribution of aggregated OOF predicted probabilities from the Elastic Net Logistic model. Samples in the high-Braak group tended to show higher predicted probabilities than those in the non-high-Braak group, while the two distributions remained partially overlapping (Figure 5c). Taken together, these results indicate that the branch-associated candidate panel contained information relevant to the prediction of high-Braak group status and achieved moderate internal predictive performance within the present cohort.

### 3.5 Lorentz t-SNE visualization

As shown in Figure 6, the standard Euclidean and Lorentz t-SNE embeddings displayed different visual organizations of the samples. In the standard Euclidean t-SNE embedding (Figure 6a), high-Braak and non-high-Braak samples were broadly intermixed across the embedded space. As an additional control, standard Euclidean t-SNE was also applied to a sample-wise Euclidean distance matrix derived from the transcriptomic expression profiles. This Euclidean-distance-based visualization also showed broad intermixing of high-Braak and non-high-Braak samples, without a clearly separated high-Braak cluster (Figure 8 in Appendix 3). In contrast, the Lorentz t-SNE embedding (Figure 6b) showed a more localized arrangement of high-Braak samples, with a subset of high-Braak samples concentrated along a branch-like region of the hyperbolic embedding.

**Figure 6:**
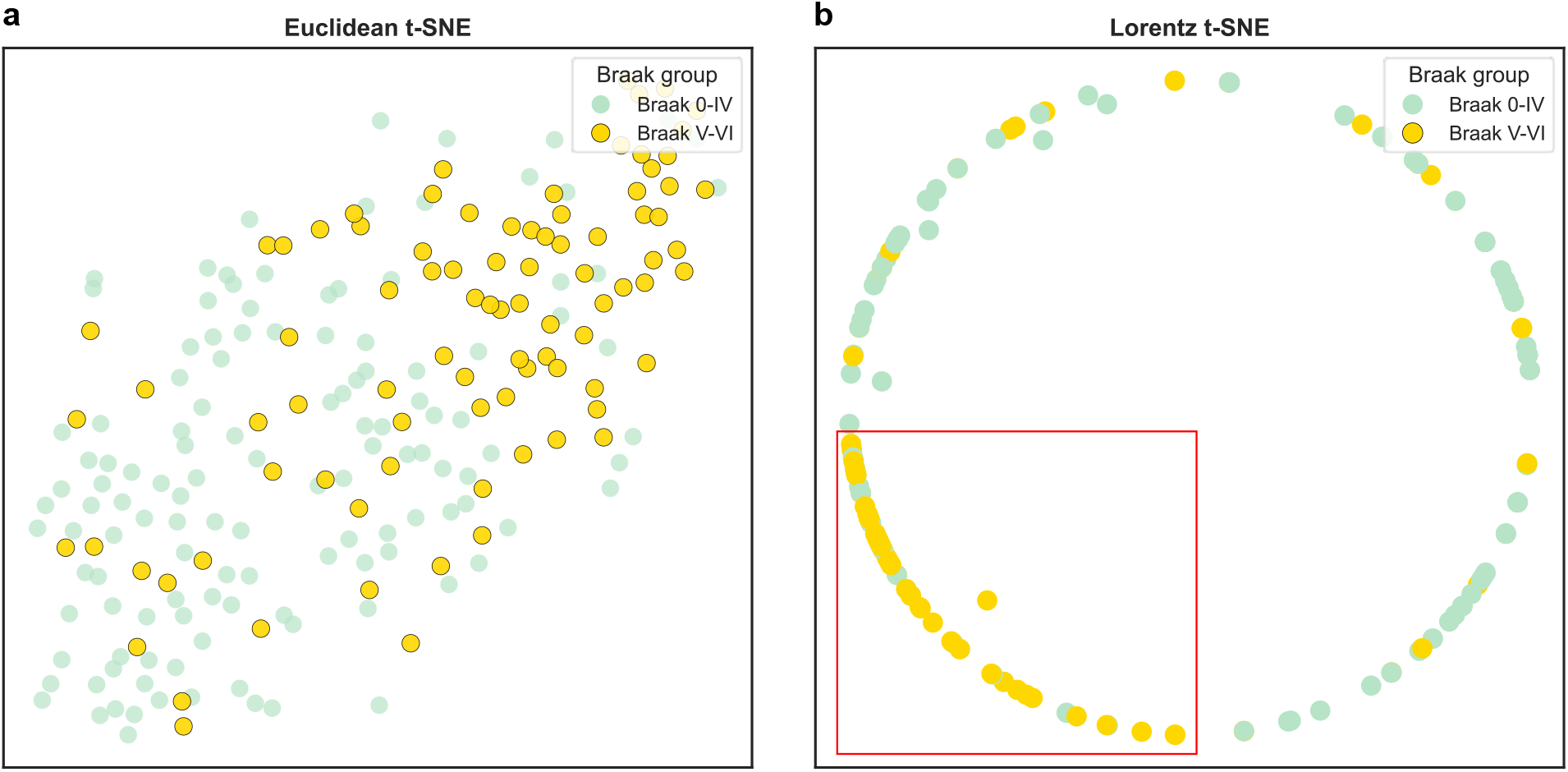
Two-dimensional visualization of MSBB BM36 samples based on the Lorentz distance matrix. **(a)** Standard Euclidean t-SNE embedding. **(b)** Lorentz t-SNE embedding optimized on ℋ^2^ and displayed in two-dimensional disk coordinates. Each point represents one sample, and colors indicate Braak-defined group status. The red rectangle highlights the local region enriched for high-Braak samples in the Lorentz t-SNE visualization.

**Figure 7:**
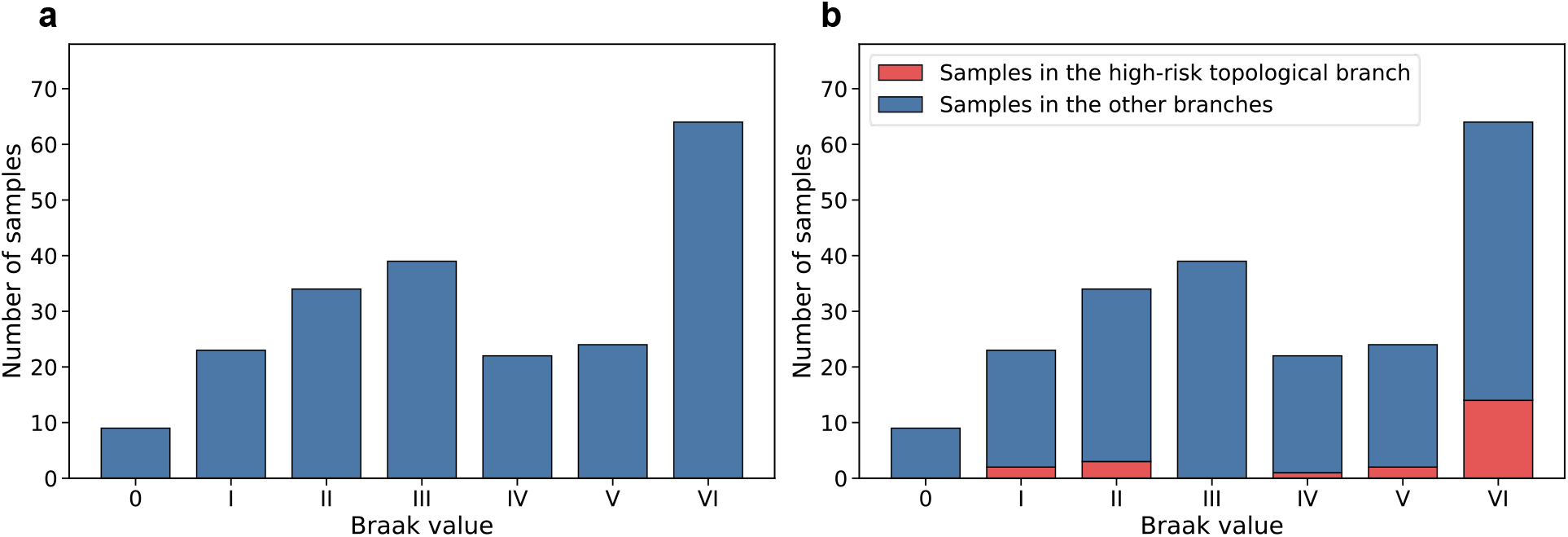
The distribution of Braak staging scores in the MSBB BM36 cohort analyzed in this study. **(a)** Overall distribution of Braak staging scores across the entire MSBB BM36 cohort. **(b)** Distribution of Braak staging scores stratified by the high-risk topological branch and all remaining branches in the MSBB BM36 cohort.

**Figure 8:**
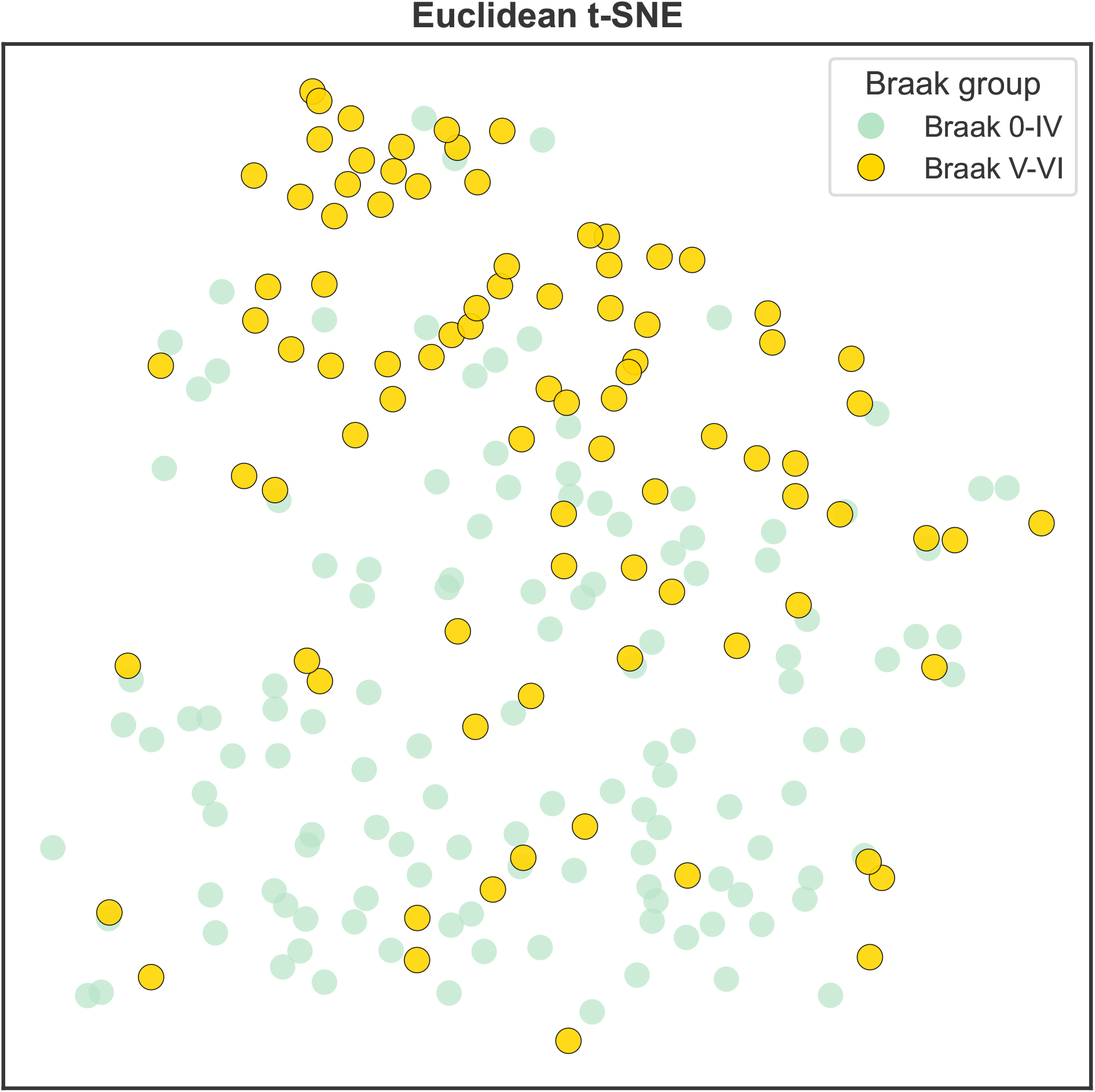
Euclidean t-SNE visualization based on the Euclidean distance matrix. Each point represents an MSBB BM36 sample, and colors indicate Braak-defined group status.

**Figure 9:**
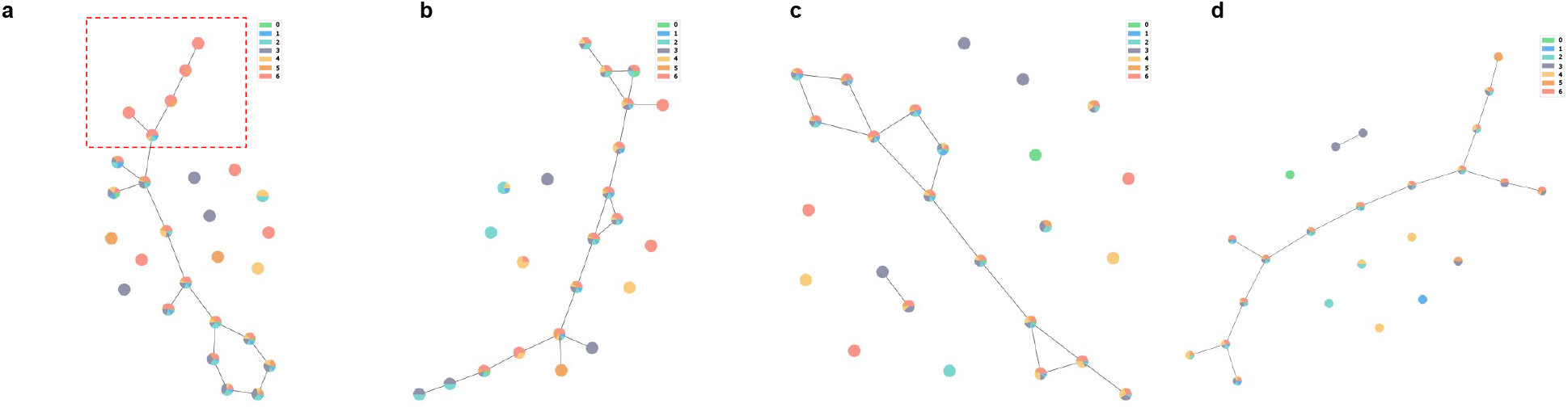
The GMM soft Mapper output graphs across four MSBB brain regions. The same preprocessing strategy and GMM soft Mapper framework were applied to transcriptomic samples from BM36, BM10, BM22, and BM44. Each node represents a cluster of samples obtained from the Mapper construction, and the pie chart within each node shows the distribution of Braak values among samples assigned to that node. The color legend in the upper-right corner of each panel corresponds to Braak values from 0 to 6. **(a)** The GMM soft Mapper output graph for BM36. The red dashed box indicates the high-risk topological branch. **(b–d)** The GMM soft Mapper output graphs for BM10, BM22, and BM44, respectively. Comparable high-risk branch-like patterns were not observed in these three brain regions under the same analytical setting. This comparison motivated the selection of BM36 for downstream branch-associated transcriptomic and predictive analyses.

This comparison should be interpreted as an exploratory visualization rather than a formal assessment of classification performance or embedding quality. Nevertheless, the observed difference is consistent with the motivation for using a hyperbolic embedding space: the GMM soft Mapper-based analysis suggested a branching organization in the MSBB BM36 transcriptomic samples, and hyperbolic geometry provides a natural representation space for data with hierarchical or tree-like organization. By defining the low-dimensional embedding on the Lorentz hyperboloid and constructing low-dimensional similarities from Lorentz geodesic distances, Lorentz t-SNE provided a visualization that was aligned with the Lorentz distance structure used in the preceding topological analysis. These results suggest that Lorentz t-SNE may complement Mapper-based analysis as an exploratory visualization tool for examining phenotype-related local organization in transcriptomic data.

## 4 Discussion

In this study, we investigated the transcriptomic features and geometry-aware visualization of a previously identified high-risk topological branch in the MSBB BM36 cohort, with a particular focus on its relationship to high-Braak group status. By integrating branch-informed co-expression network analysis, repeated stability-based differential expression analysis, functional enrichment, and nested cross-validation, we identified a branch-associated candidate gene panel that contained predictive information related to high-Braak group status within the present cohort. In parallel, our introduced Lorentz t-SNE provided an exploratory hyperbolic visualization of the sample-wise Lorentz distance structure, in which high-Braak samples showed a more localized arrangement than in the corresponding Euclidean t-SNE embedding. Taken together, these findings suggest that the Mapper-derived high-risk branch may capture transcriptomic variation associated with Braak-defined grouping, rather than representing only an abstract topological partition.

A central aspect of this study is that its feature discovery strategy was organized around the topology-derived branch rather than directly around the high-Braak prediction target. RNA-sequencing-based studies have reported molecular subtypes of Alzheimer’s disease defined from transcriptomic profiles across large postmortem brain cohorts [Neff et al., 2021]. These subtype-oriented analyses primarily aim to define molecular subgroups of Alzheimer’s disease, whereas this study focuses on the transcriptomic and predictive relevance of a Mapper-derived high-risk topological branch. Therefore, the high-risk topological branch examined here should be interpreted as a topology-derived organization of transcriptomic samples rather than as a conventional transcriptomic subtype. The WGCNA hub-gene selection and repeated differential expression analysis were both branch-associated, and the resulting candidate panel was subsequently evaluated for its relationship to high-Braak group status using a nested cross-validation framework. This design does not eliminate the need for external validation, but it helps reduce the risk that the predictive analysis simply reflects direct feature selection on the same group label used for model evaluation. The moderate internal predictive performance observed across repeated outer-fold evaluations therefore supports the presence of branch-associated transcriptional information related to Braak-defined group status, while remaining insufficient to establish a clinically generalizable biomarker model.

The Lorentz t-SNE analysis further provided a geometry-aware visualization of the same sample organization. Because the high-risk branch was originally identified from a Lorentz distance-based topological analysis, embedding the samples on the Lorentz hyperboloid offered a way to examine whether the corresponding distance structure could be represented in a low-dimensional hyperbolic space. The observed local aggregation of high-Braak samples in the Lorentz t-SNE embedding is compatible with the possibility that part of the AD-related transcriptomic variation in this cohort may have a branch-like or hierarchical organization. However, this result should be interpreted as qualitative and exploratory. The present analysis does not establish that the full transcriptomic manifold is intrinsically hyperbolic, nor does it provide a formal benchmark demonstrating superiority over Euclidean embeddings across datasets. Instead, the proposed Lorentz t-SNE should be viewed here as a complementary visualization tool for examining topology-derived sample organization.

Several limitations should be acknowledged. First, the study was conducted in a single brain region from a single cohort, with a modest number of samples and a small high-risk topological branch. Independent validation in additional AD transcriptomic cohorts, other MSBB brain regions, or harmonized multi-cohort datasets will be required to assess the generalizability of the branch-associated gene panel and the reduced Elastic Net model. Second, the prediction task focused on cross-sectional Braak-defined group status rather than longitudinal disease progression or clinical conversion; therefore, the model should not be interpreted as a prognostic or diagnostic tool. Third, although nested cross-validation was used for internal model evaluation, the candidate feature panel was defined before model training based on branch-associated stability selection and network filtering. Accordingly, the reported performance estimates should be interpreted as internal evaluation of a predefined branch-associated feature set, not as validation of a fully independent feature-discovery pipeline. Fourth, the Lorentz t-SNE results were assessed visually and were not accompanied by quantitative measures of neighborhood preservation, embedding stability, or cross-dataset reproducibility. Finally, the biological roles of the retained genes and enriched pathways require further experimental or independent transcriptomic validation.

Despite these limitations, this study illustrates a topology-guided application framework for characterizing AD-related transcriptomic heterogeneity. By connecting the GMM soft Mapper-derived high-risk branch with the co-expression modules, stable branch-associated genes, internal predictive modeling, and Lorentz-hyperbolic visualization, the analysis suggests that topology-derived sample structures may reflect biologically relevant variation in AD brain transcriptomes. Future work should evaluate whether similar topology-guided and geometry-aware strategies can complement the interpretation of transcriptomic heterogeneity across additional AD cohorts, brain regions, disease stages, and multi-omics datasets. Moreover, future studies could extend this analysis to transcriptomic datasets from brain regions in mice and other relevant organisms that have corresponding or anatomically related structures, while also incorporating empirical validation in mouse models. This would require appropriate cross-species orthology mapping and careful evaluation of anatomical correspondence across brain regions and species. Such cross-species integrative analyses, combined with experimental validation, may help determine whether the branch-associated genes and enriched biological processes identified in the present study display conserved or comparable disease-related patterns across species and experimental systems, thereby further enhancing the biological interpretability and translational relevance of these findings.

## 5 Data availability

The paper employs publicly accessible datasets in experiments. The RNA expression sequences used in this manuscript are obtained from the Mount Sinai/JJ Peters VA Medical Center Brain Bank [Wang et al., 2018] and are available at https://www.synapse.org/ with SynID 20801188. The results of all experiments, and the code are available at: https://github.com/zchhh01/HyperIAD.

## 6 Acknowledgments

The authors thank ShanghaiTech University for supporting this work through the startup fund and the HPC platform.

## Appendix 1

### Geometry of the Lorentz hyperboloid

Hyperbolic space admits several representations, notably the Lorentz hyperboloid and Poincaré ball models [Ratcliffe, 2006, Anderson, 2005]. The Lorentz hyperboloid model is particularly convenient for optimization in hyperbolic representation learning [Nickel and Kiela, 2018, Mishne et al., 2023]. In this work, the low-dimensional embedding is formulated on the forward sheet of the Lorentz hyperboloid in Minkowski space. Let **u, v** ∈ℝ^*n*+1^ be vectors written as **u** = (*u*_0_, *u*_1_, …, *u*_*n*_) and **v** = (*v*_0_, *v*_1_, …, *v*_*n*_). Their Lorentzian inner product is defined by

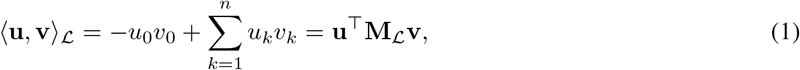

where

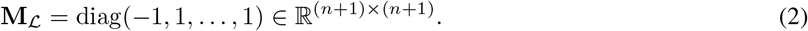

Under this bilinear form, the Lorentz hyperboloid is given by

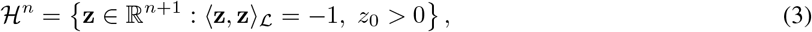

which corresponds to the forward sheet of a two-sheeted hyperboloid.

For two points **z**_*i*_, **z**_*j*_ ∈ ℋ^*n*^, the geodesic distance induced by the Lorentz metric is

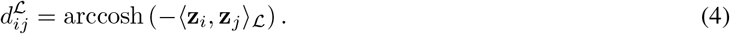

The tangent space at a point **z** ∈ ℋ^*n*^ is

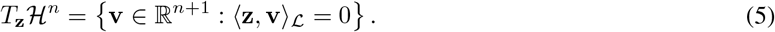

For any tangent vector **v** ∈ *T*_**z**_ℋ^*n*^, its Lorentz norm is defined as

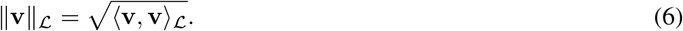

This tangent-space structure provides the geometric setting in which gradient-based updates are later carried out on the manifold.

Although optimization in this study is carried out in the Lorentz model, it is useful to note its equivalence to the Poincaré ball model [Ratcliffe, 2006, Anderson, 2005, Nickel and Kiela, 2018]. To make this correspondence explicit, let

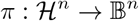

denote the mapping from the Lorentz hyperboloid to the Poincaré ball, and let

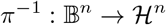

denote its inverse. Then, for **z** = (*z*_0_, **z**^*′*^) ∈ ℋ ^*n*^, the associated Poincaré coordinate is given by

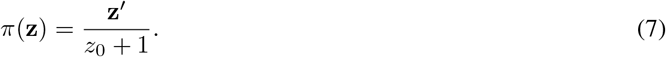

Conversely, for **u** ∈ B^*n*^ with ∥**u**∥ < 1, the corresponding point on the Lorentz hyperboloid is

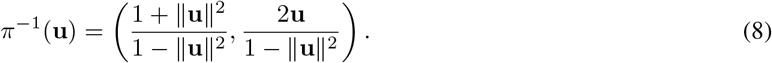

These standard coordinate transformations show that the Lorentz and Poincaré models are isometric representations of the same hyperbolic space, while the Lorentz representation remains convenient for the optimization procedure adopted here.

### Optimization on the Lorentz hyperboloid

For completeness, we first recall the standard t-SNE formulation [Maaten and Hinton, 2008] before introducing its Lorentz-hyperbolic counterpart. Given high-dimensional observations {**x**_1_, …, **x**_*N*_ } ⊂ℝ^*d*^, the conditional similarities are defined as

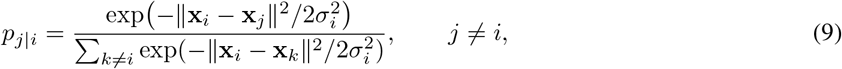

and the symmetric joint probabilities are given by

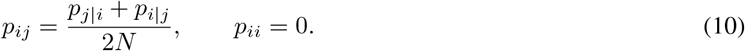

In the standard Euclidean setting, the low-dimensional similarities are constructed as

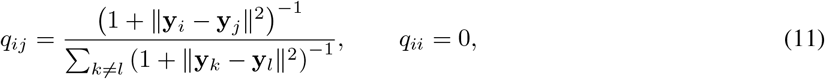

and the embedding is obtained by minimizing the Kullback–Leibler divergence

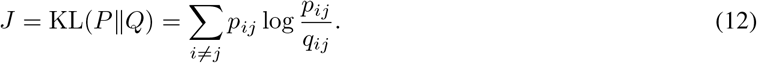

In the Lorentz t-SNE formulation adopted in this study, the Euclidean embedding space is replaced by the Lorentz hyperboloid. The embedded points are denoted by **z**_*i*_ ∈ ℋ^*n*^, and their low-dimensional affinities are defined through Lorentz geodesic distances. Specifically, for *i* ≠*j*,

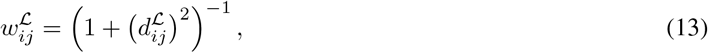

where 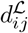 denotes the Lorentz geodesic distance between **z**_*i*_ and **z**_*j*_. The normalized Lorentz low-dimensional similarities are

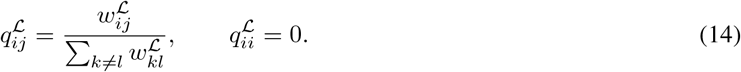

Accordingly, the Lorentz-hyperbolic objective function is

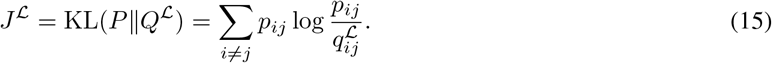

The optimization is carried out on the Lorentz hyperboloid rather than in a linear Euclidean space. Therefore, the Euclidean coordinate gradient in the ambient space must be converted according to the Lorentzian metric and projected onto the tangent space of the hyperboloid. The derivation of the tangent-space gradient used for the update is given in the next subsection.

To keep each update on the hyperboloid, the tangent-space update is mapped back to the manifold through the exponential map, as in standard Riemannian optimization [Absil et al., 2008, Bonnabel, 2013]:

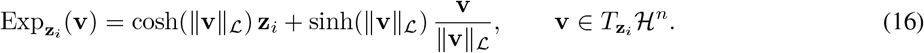

When ∥**v**∥_*ℒ*_ = 0, the exponential map is understood by continuity as 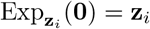. Let *η* denote the learning rate. Using the tangent-space gradient defined below, one gradient-based update is written as

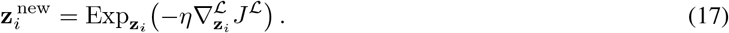

### Derivation of the Lorentz t-SNE Gradient

Throughout this subsection, 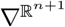 denotes differentiation with respect to the ambient Euclidean coordinates. The manifold ℋ^*n*^ denotes the Lorentz hyperboloid, whereas the superscript ℒ is used for quantities defined with respect to the Lorentzian metric. In particular, 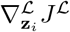 denotes the tangent-space gradient obtained after metric conversion and projection onto 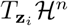.

This subsection derives the Lorentz-hyperbolic gradient used in the update rule from the ordinary Euclidean coordinate gradient in the ambient space, following the standard Riemannian optimization principle of converting ambient derivatives into tangent-space gradients [Absil et al., 2008].

Here 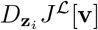 denotes the first-order variation of the objective when only the *i*-th embedded point is perturbed in the direction **v**. Equivalently,

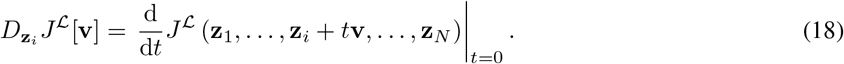

For the Lorentz t-SNE objective *J* ^*ℒ*^, the Euclidean coordinate gradient with respect to the *i*-th embedded point is characterized by the partial directional derivative

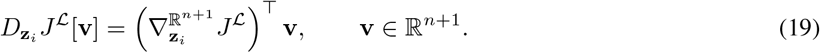

On the Lorentz hyperboloid, ambient vectors are paired through the Lorentzian inner product

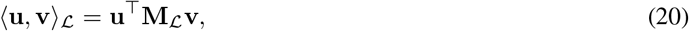

where **M**_**ℒ**_ is the Lorentz metric matrix defined above. The same differential can therefore be represented under the Lorentzian inner product as

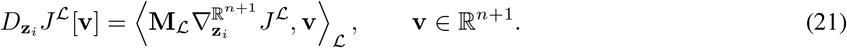

Indeed,

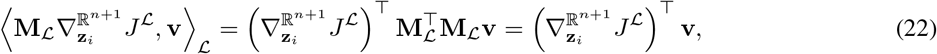

since 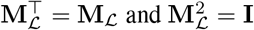.

The resulting ambient vector is not necessarily tangent to the hyperboloid. Since the tangent space at **z**_*i*_ ∈ ℋ^*n*^ is

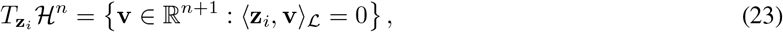

the Lorentz-orthogonal projection onto 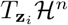 gives

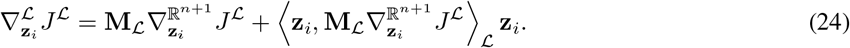

This vector is tangent because

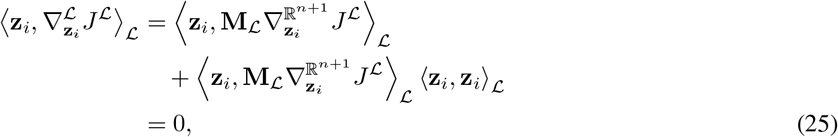

as ⟨**z**_*i*_, **z**_*i*_⟩_ℒ_ = −1 on ℋ^*n*^.

It remains to compute the Euclidean coordinate gradient 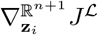. The Lorentz geodesic distance is

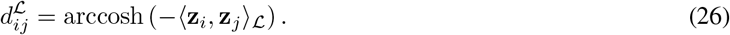

For notational clarity, define

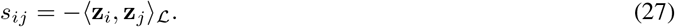

For *i* ≠ *j* and non-coincident embedded points, *s*_*ij*_ > 1, so the derivative of arccosh is well defined. Since

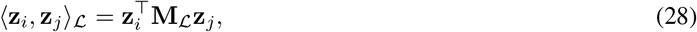

the Euclidean coordinate derivative of *s*_*ij*_ with respect to **z**_*i*_ is

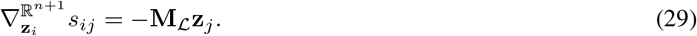

Together with

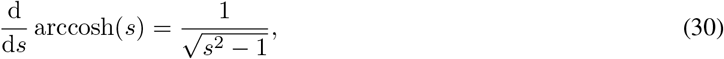

#### Algorithm 1 Riemannian gradient descent step for Lorentz t-SNE

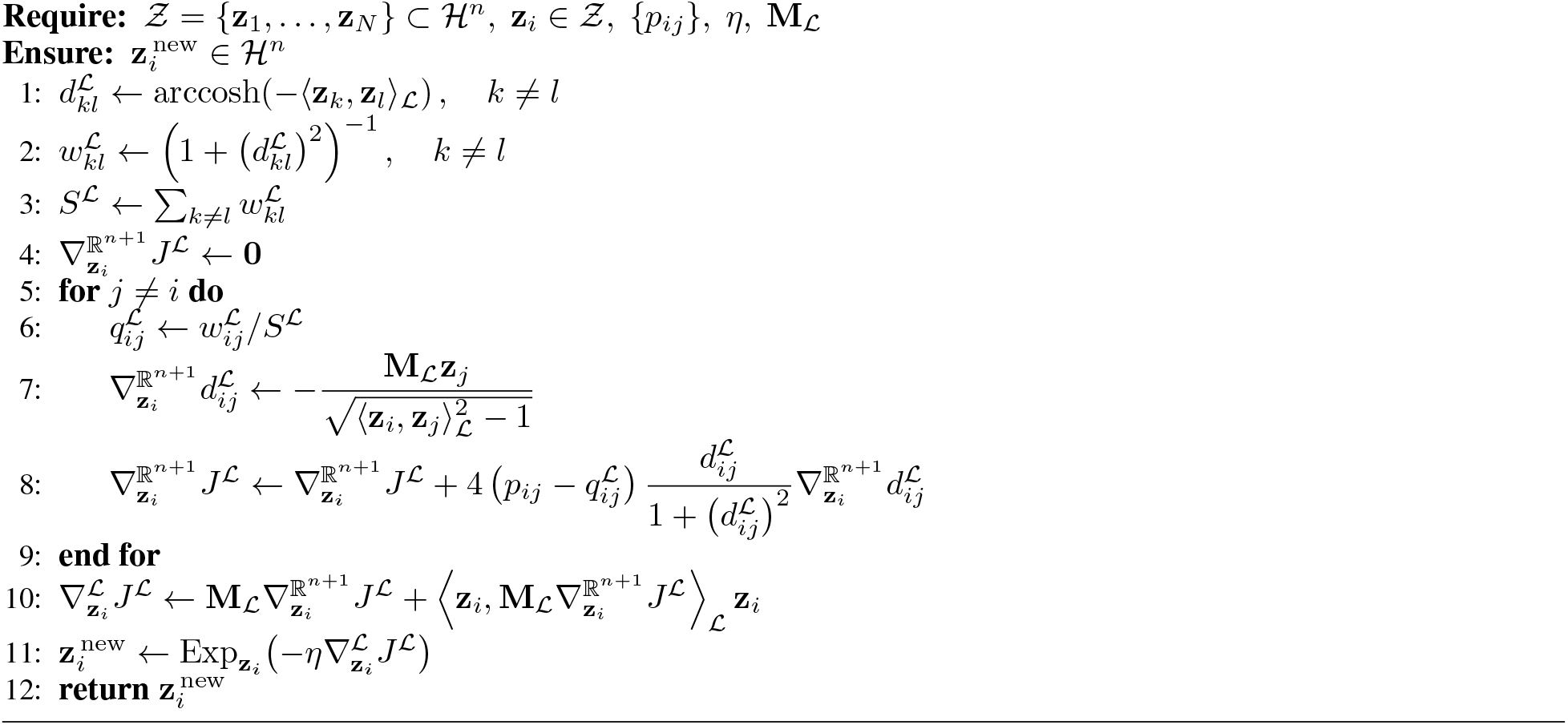

this yields

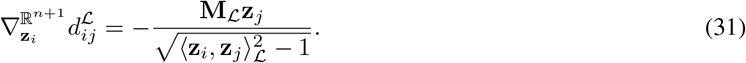

The low-dimensional Lorentz affinity is based on

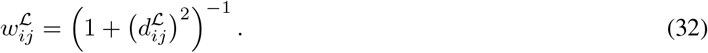

With

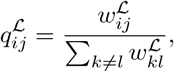

together with the symmetry of *P* and *Q*^*ℒ*^, differentiation of the Kullback–Leibler objective gives

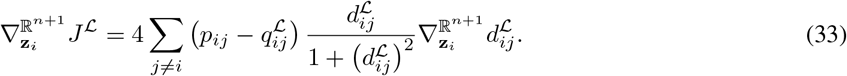

Substituting the Euclidean coordinate derivative of the Lorentz distance gives the expanded expression

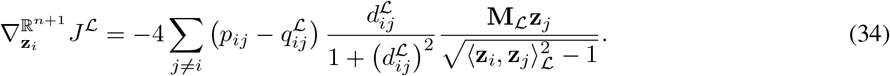

Finally, the tangent-space gradient used in the Lorentz-hyperbolic update is obtained by substituting this Euclidean coordinate gradient into the metric conversion and projection formula:

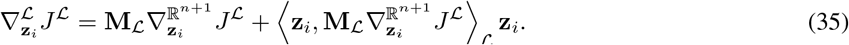

This tangent vector is the quantity used in the exponential-map update in the previous subsection.

## Appendix 2

### Sample composition and Braak-value distribution

To provide a visual summary of the study cohort, we examined the distribution of Braak values and the relationship between Braak-defined grouping and topology-derived branch membership. The final dataset contained 215 samples, including 88 high-Braak samples (Braak V–VI) and 127 non-high-Braak samples (Braak 0–IV). Figure 7 summarizes the Braak-value distribution in the full BM36 dataset and after stratification by high-risk topological branch membership.

### Model coefficients and standardization parameters

For the full-cohort refitted Elastic Net logistic regression model, input expression values were standardized before prediction. For each retained gene, the standardized expression value was calculated as

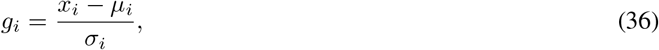

where *x*_*i*_ denotes the processed expression value of the *i*th gene, and *µ*_*i*_ and *σ*_*i*_ denote the mean and standard deviation used for feature scaling, respectively. The fitted model was then represented as the following linear predictor:

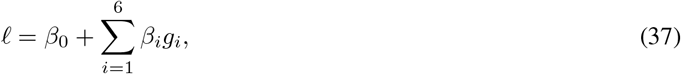

where *β*_0_ is the intercept, *β*_*i*_ is the Elastic Net coefficient of the *i*th retained gene, and *g*_*i*_ is the standardized expression value of that gene. The predicted probability of high-Braak group membership was obtained using the logistic transformation:

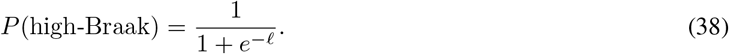

The coefficients and scaling parameters of the retained genes in the full-cohort refitted Elastic Net model are listed in Table 2. These parameters were used to describe the refitted model structure, whereas predictive performance was evaluated using held-out outer-fold predictions generated during nested cross-validation.

## Appendix 3

### Euclidean t-SNE visualization using a Euclidean distance matrix

The t-SNE control analysis using the Euclidean distance matrix is shown in Figure 8.

### The GMM soft Mapper outputs on the four brain regions

To provide a supplementary comparison across brain regions, Appendix Figure 9 shows the GMM soft Mapper output graphs generated separately for BM36, BM10, BM22, and BM44 using the same preprocessing and Mapper construction framework.

